# Noise improves the association between effects of local stimulation and structural degree of brain networks

**DOI:** 10.1101/2023.01.10.523529

**Authors:** Yi Zheng, Shaoting Tang, Hongwei Zheng, Xin Wang, Longzhao Liu, Yaqian Yang, Yi Zhen, Zhiming Zheng

**Affiliations:** School of Mathematical Sciences, Beihang University, Beijing, China; Institute of Artificial Intelligence, Beihang University, Beijing, China; Key laboratory of Mathematics, Informatics and Behavioral Semantics (LMIB), Beihang University, Beijing, China; State Key Lab of Software Development Environment (NLSDE), Beihang University, Beijing, China; Zhongguancun Laboratory, Beijing, P.R.China; Beijing Advanced Innovation Center for Future Blockchain and Privacy Computing, Beihang University, Beijing, China; PengCheng Laboratory, Shenzhen, China; Institute of Medical Artificial Intelligence, Binzhou Medical University, Yantai, China; School of Mathematical Sciences, Dalian University of Technology, Dalian, China; Beijing Academy of Blockchain and Edge Computing (BABEC), Beijing, China

## Abstract

Stimulation to local areas remarkably affects brain activity patterns, which can be exploited to investigate neural bases of cognitive function and modify pathological brain statuses. There has been growing interest in exploring the fundamental action mechanisms of local stimulation. Nevertheless, how noise amplitude, an essential element in neural dynamics, influences stimulation-induced brain states remains unknown. Here, we systematically examine the effects of local stimulation by using a large-scale biophysical model under different combinations of noise amplitudes and stimulation sites. We demonstrate that noise amplitude nonlinearly and heterogeneously tunes the stimulation effects from both regional and network perspectives. Furthermore, by incorporating the role of the anatomical network, we show that the peak frequencies of unstimulated areas at different stimulation sites averaged across noise amplitudes are highly positively related to structural connectivity. Crucially, the association between the overall changes in functional connectivity as well as the alterations in the constraints imposed by structural connectivity with the structural degree of stimulation sites is nonmonotonically influenced by the noise amplitude, with the association increasing in specific noise amplitude ranges. Moreover, the impacts of local stimulation of cognitive systems depend on the complex interplay between the noise amplitude and average structural degree. Overall, this work provides theoretical insights into how noise amplitude and network structure jointly modulate brain dynamics during stimulation and introduces possibilities for better predicting and controlling stimulation outcomes.

**Author summary:** Despite the extensive application of local stimulation in cognition research and disease treatments, how regional perturbations alter brain-wide dynamics has not yet been fully understood. Given that noninvasive stimulation is associated with changes in the signal-noise relationship, we assume that noise amplitude is one of the plausible factors modulating the stimulation effects. Using a whole-brain biophysical model under different stimulation sites and noise amplitudes, we explore the influence of noise amplitude on stimulation effects and, more importantly, the interplay between noise amplitude and network structure. From a regional perspective, noise amplitude reduces the peak frequencies in unstimulated areas during stimulation. Moreover, we find a high similarity between the noise-averaged peak frequency matrix and the structural network. From a network perspective, we show that the changes in functional connectivity are decreased by noise amplitude, while the alterations in structural constraints display nonmonotonic trends. Intriguingly, increasing the noise amplitude in specific ranges can improve the association between network-level effects and structural degree, promoting better predicting and controlling therapeutic performance. Finally, the behaviors of cognitive systems quantified by network-level effects are jointly modulated by the noise amplitude and average structural degree.

## Introduction

Complex interactions among brain areas elicit the rich spatiotemporal profiles of neural activity that underlie human cognition and behaviors [1, 2]. Because the brain is an open system, typical brain activity patterns are highly influenced by local perturbations, yielding various dynamical states [3]. For example, sensory inputs can be viewed as local stimulation, which may lead to neural activity changes in primary areas and affect the associative cortex through neural circuits, thus supporting sophisticated cognitive processes such as learning, decision-making and memory [4–6]. In addition, aberrant statuses caused by certain brain disorders are associated with local perturbations. Specifically, some generalized epileptic seizures are caused by stimulus-induced abnormal activity in focal areas spreading throughout the brain [7, 8]. Despite the critical role of local perturbations, how regional stimulation modulates the underlying neural processes has not yet been fully established [9].

Since their inception, artificial stimulation techniques have served as efficient tools that allow researchers to directly investigate responses to experimentally altered local neural activity. These methods have been widely used to explore the causal relationship between select brain regions and cognitive processes or task behaviors [10]. Importantly, they are also promising in clinical applications for the treatment of psychiatric and neurological disorders. For example, deep brain stimulation (DBS) is commonly used for patients with Parkinson’s disease, Alzheimer’s disease and dementia [11–13]. Moreover, transcranial magnetic stimulation (TMS) is often employed for treating epilepsy, autism and schizophrenia [14–16]. Revealing the effects of local stimulation may improve our understanding of the neurodynamic bases of human cognition and behaviors and facilitate the development and utilization of stimulation techniques.

It has been widely accepted that local perturbations not only induce regional modifications near stimulation sites but also provoke broad system-level impacts [10, 17, 18]. Since neural activity propagates along white matter bundles, researchers have explored how anatomical connectivity constrains global stimulation effects, highlighting the critical contribution of macroscale structural properties such as degree and modularity [19, 20]. Moreover, recent studies have demonstrated that stimulation effects rely on physiological and cognitive states [21, 22]. Compared to the resting state, sleep or working memory states generate specific neural activity patterns that alter the transmission of local stimulation [23, 24], leading to differences in the regional power spectra, interregional functional coupling, and behavioral performance. Despite these advances, our poor understanding of the high variability of stimulation outcomes across subjects suggests that the fundamental mechanisms that explain how regional activity changes alter brain-wide dynamics need to be studied further [25, 26]. The activity patterns elicited by stimulation should be jointly modulated by multiple neurophysiological factors. For example, recent research showed that the response to local perturbations depends on both the stimulation sites and oscillatory states of brain network activity [27]. Nevertheless, most studies have tended to examine the impact of single elements, thus overlooking other factors that may have vital influences on network communication and ignoring the essential interplay among these factors.

Neural noise, including multiple sources such as sensory, cellular and electrical noise, affects all aspects of the behaviors of the nervous system [28]. On the one hand, neural noise is thought to hinder information processing and transmission. On the other hand, neural noise has been found to help maintain and promote brain function, including shaping resting-state functional networks [29, 30], enhancing neural synchronization [31, 32] and affecting task performance [33, 34]. In addition, previous research have indicated that the brain, as a noisy dynamical system, manifests subject-specific parameters at various scales, thus producing diverse outputs [28, 35]. Moreover, a recent study related local stimulation to noise amplitude by showing that the impact of noninvasive brain stimulation could be viewed as a neural activity modification that alters the signal-noise relationship [36]. Based on this evidence, we assume that neural noise is a crucial factor that influences stimulation-induced brain states. However, how the noise amplitude is related to the global consequences of local stimulation remains unknown. In particular, despite the previously discovered significant contribution of network structure, the interplay between noise amplitude and network structure remains unexplored.

Experimentally examining the effects of local stimulation across different parameters is impractical, time-consuming and potentially detrimental to participants; however, model-based numerical simulation offers a powerful approach to investigating these unknown situations [27, 37–41]. Thus, in the present work, we utilize a whole-brain biophysical model composed of Wilson-Cowan neural masses to systematically explore dynamic brain states at different stimulation sites under various noise amplitudes. We first choose the appropriate global coupling strength, which is independent of the noise amplitude, before stimulation and then evaluate the stimulation effects by examining the frequencies associated with the maximum values in the regional power spectrum (peak frequencies), the changes in the functional configurations (functional effects) and the alterations in the structural constraints on function (structural effects) [37].

From the regional perspective, we show that the noise amplitude influences the peak frequencies of unstimulated brain areas, shifting the frequencies from higher to lower values. Moreover, we find a high positive association between the peak frequencies of unstimulated areas at different stimulation sites averaged across noise amplitudes and the corresponding structural connectivity, underlining the antagonistic effects of the direct connection strength and noise amplitude. From the network perspective, we show that functional effects are nonlinearly weakened by noise amplitude, while structural effects exhibit nonmonotonic trends. Importantly, due to the heterogeneous role of noise amplitude on stimulation sites, increasing the noise amplitude in specific ranges can enhance both the Pearson correlation coefficient and the adjusted coefficient of determination between the functional or structural effects and the structural degree of stimulation sites, potentially improving prediction and control in clinical intervention approaches. The changes in the noise amplitude can even turn the correlation of structural effects from negative to positive. Finally, we show that the noise amplitude and system-level average degree jointly modulate the performance of cognitive systems in terms of functional and structural effects. The subcortical system with a high average degree exhibits distinct behaviors under various noise amplitudes from the sensory and association system with a low average degree. In summary, our study highlights the importance of the coupling between noise amplitude and network structure in influencing the effects of regional stimulation, thereby providing further insights into the fundamental principles of brain dynamics and contributing to the development of personalized stimulation techniques and the optimization of therapeutic performance.

## Results

We utilize a three-step investigation procedure that consists of Input data, Computational model, and Analysis scheme (Fig. 1); the details of this approach are provided in the Materials and methods. Specifically, we first present the structural network, distance network, and stimulation protocol as the main inputs of the procedure. The group-level structural connectivity matrix (Fig. 1A) is estimated according to the diffusion-weighted MRI of 30 healthy subjects, combined with a parcellation of 82 regions (including subcortical areas) following the atlas [42]. The matrix elements are fixed as the number of fibers between brain regions normalized by the geometric mean of their volumes and capture the strength of interactions between brain areas to some extent. The group-representative distance matrix (Fig. 1B) is derived as the mean Euclidean distance between the centers of brain areas across subjects. This matrix is used to estimate the time delays associated with interareal communication. External stimulation with an intensity of 1.25 is applied to the brain during the third second but is absent for the first two seconds (Fig. 1C).

**Fig 1.**
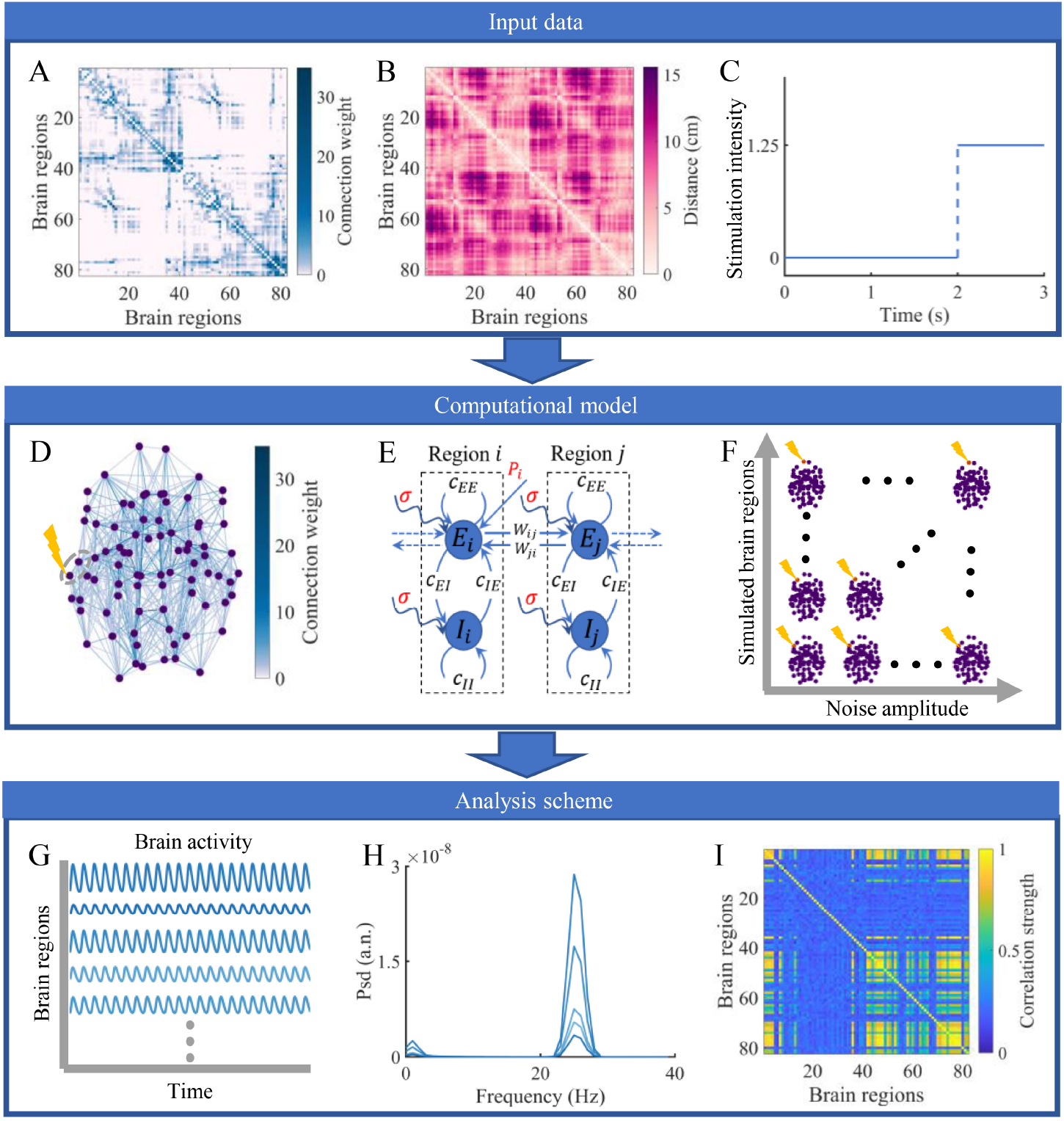
Workflow consisting of Input data, Computational model and Analysis scheme. (A) Group-level structural connectivity matrix based on 82-node brain parcellation. (B) Group-averaged distance matrix characterized by the same parcellation. (C) External stimulation protocol, with an intensity of 0 for the first two seconds and an intensity of 1.25 for the third second. (D) An example of local stimulation in the structural brain network. The purple dots and blue lines represent the centers of the brain regions and the strongest 20% of connections between them. The line darkness is positively related to the connection weight. The stimulated brain region indicated by the yellow lightning bolt and its unstimulated neighbor are circled to demonstrate the dynamics. (E) Schematic of two brain regions with Wilson-Cowan dynamics linked by excitatory connections. Each region includes coupled excitatory and inhibitory populations disturbed by noise with amplitude *σ*. An external perturbation *P*_*i*_ is applied to region i to increase the excitatory input. (F) Simulation experimental design. Different brain regions are stimulated under various noise amplitudes. (G) Time series of excitatory activity for each brain region generated by the computational model. (H) Power spectrum estimation to quantify the stimulation outcomes from a regional perspective. (I) Functional connectivity matrix used to evaluate network-level effects.

A single trial involving local perturbation to the brain network is illustrated in Fig. 1D. The stimulation, which is indicated by the yellow lightning bolt, is applied to a single area. The stimulated area and one of its neighboring areas are surrounded by a gray dotted line, and their dynamics are shown in Fig. 1E. A Wilson-Cowan neural mass composed of coupled excitatory and inhibitory populations is inserted into each region [43]. Both neural populations are disturbed by noise with a specific amplitude. The local perturbation is assumed to act on the excitatory population in the stimulated region. Interareal communication is achieved through long-range structural connections linking excitatory populations in different regions [27, 37, 44]. To systematically explore the role of noise amplitude in modifying the stimulation outcomes, we perform simulations under different combinations of noise amplitudes and stimulation sites (Fig. 1F), each with 30 realizations.

After the simulations, we extract the firing rates of all excitatory populations during 1-2 s and 2-3 s as time series before and during stimulation to evaluate the impacts of the perturbations (Fig. 1G). First, from a regional perspective, one crucial effect of local stimulation is altering the oscillations in different brain areas, which are commonly related to brain functions and behaviors [45, 46]. Therefore, we used the characteristics of the power spectrum during stimulation, such as the frequency corresponding to the maximum power (peak frequency) to reflect the effects of stimulation propagation (Fig. 1H). Second, the dynamic information of the brain is stored not only in individual regions but also in interactions between areas. Thus, we examined the network-level stimulation effects based on the functional connectivity matrix (Fig. 1I) by quantifying the average changes in functional networks (functional effects) and the alterations in the similarity between structural and functional connectivity (structural effects) [37]. The calculation details and a summary of the measures used in this work are presented in the Materials and methods.

### The effects of the noise amplitude and global coupling strength on brain states without stimulation

To select the optimal global coupling strength *c* and provide prior knowledge about the brain state under different noise amplitudes *σ*, we perform 2-second simulations without stimulation and investigate system behaviors under various combinations of *σ* and *c*. We change *σ* from 10^*−*9^ to 10^*−*2^ and *c* from 0.01 to 0.3.

In Fig. 2A, we present the time-averaged excitatory activity 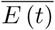 as a function of the global coupling strength at a single noise amplitude (*σ* = 10^*−*5^). The results show that there exists a threshold of c, above which the 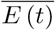 values in most regions change sharply. Fig. 2B shows how the time- and network-averaged activity 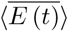 varies with the noise amplitude and global coupling strength. The threshold remains constant and is unrelated to the noise amplitude. We next examine the network-averaged peak frequency and peak power (⟨*f*_*peak*_)*and*(*p*_*peak*_⟩). We find the same threshold in Fig. 2C and Fig. 2D. Below this threshold, the results show that ⟨*f*_*peak*_⟩ is less than 10 HZ and ⟨*p*_*peak*_⟩ is relatively low. Note that a larger noise amplitude leads to a larger ⟨*p*_*peak*_⟩. Moreover, slightly above the threshold, ⟨*f*_*peak*_⟨ and ⟨*p*_*peak*_⟨ are both considerably enhanced. These results are consistent with those in previous studies, indicating that crossing the threshold causes most nodes in the system to undergo bifurcations from low-activity steady states with fluctuations mainly driven by noise to high-amplitude oscillatory states [27, 37, 41].

**Fig 2.**
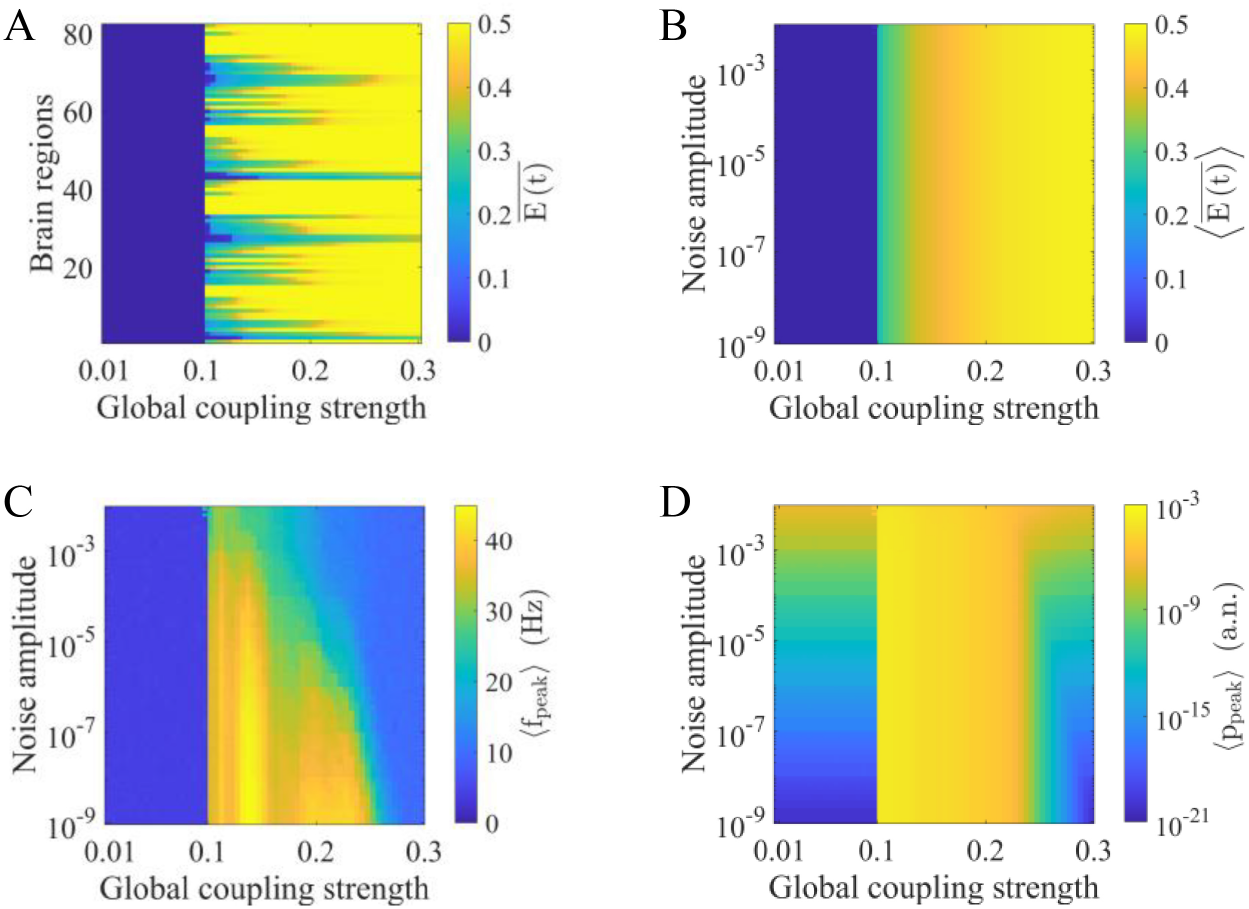
The effects of noise amplitude and the global coupling strength on brain states before stimulation. (A) Time-averaged excitatory activity 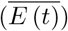 in each brain region under different global coupling strengths when *σ* = 10^*−*5^. (B) Time- and network-averaged excitatory activity 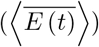 under different combinations of global coupling strengths and noise amplitudes. (C) Network-averaged peak frequency of the excitatory activity (⟨*f*_*peak*_⟩) under different combinations of global coupling strengths and noise amplitudes. (D) Network-averaged peak power of the excitatory activity (⟨*p*_*peak*_⟩) under different combinations of global coupling strengths and noise amplitudes.

Overall, Fig. 2B-D reveals that a noise-independent threshold separates the two dynamical regimes. According to previous research [37, 41], we thus choose the value just below the threshold as the optimal global coupling strength (*c* = 0.1). This approach is consistent with the widely accepted assumption that empirical brain function is best captured by the fluctuation regime, which provides maximal flexibility in information processing [38, 47, 48]. At the chosen value, *σ* has little effect on 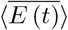 and ⟨*f*_*peak*_⟩, except for ⟨*p*_*peak*_⟩. This result indicates that the noise amplitude does not produce qualitative changes in brain states, providing a relatively uniform baseline. Note that there are several high ⟨*f*_*peak*_⟩ values at large *σ* when *c* = 0.1 in Fig. 2C. This is because large noise amplitudes combined with the initial values of simulations may result in oscillations in some regions with a limited number of realizations, which has little impact on the following analyses. We also show the distributions of ⟨*f*_*peak*_⟩ and ⟨*p*_*peak*_⟩ in all brain regions under the chosen *c* and different *σ* in S1 Fig and S2 Fig to provide additional information.

### The regional peak frequency during stimulation depends on the interplay between the noise amplitude and structural connectivity strength

In this section, we explore the peak frequency in both stimulated and unstimulated brain regions at a global coupling strength of 0.1. We mainly focus on the following problems: How does noise amplitude affect the regional peak frequency during stimulation? Moreover, given the important role of the anatomical network in shaping neural dynamics [49], how do the noise amplitude and structural properties jointly modulate the regional peak frequency?

Fig. 3A presents the peak frequencies in stimulated brain areas 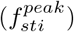 as a function of the noise amplitude and stimulation site. The external perturbation drives the stimulated region to an oscillatory state, leading to higher 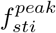 values than the case without stimulation. The results show that 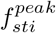 is related to the stimulation site but is rarely affected by the noise amplitude. Note that different stimulation sites influence the transmission pathways for the altered neural activity and are assumed to reflect properties of the structural brain network.

**Fig 3.**
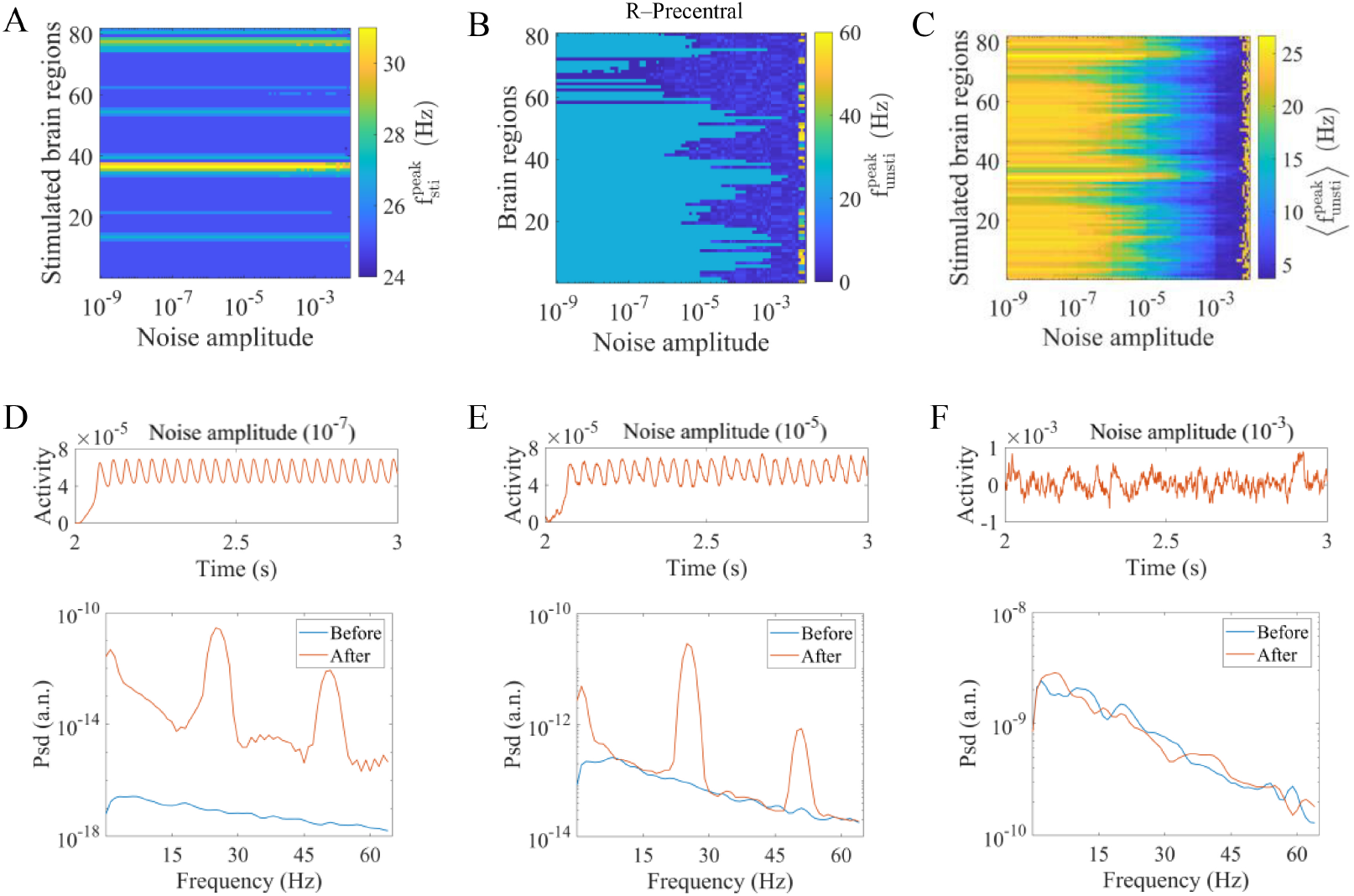
The effects of noise amplitude and the stimulation site on the regional peak frequency during stimulation. (A) Peak frequency of stimulated brain regions 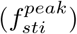 under different combinations of noise amplitudes and stimulation sites. (B) Peak frequency of unstimulated brain regions 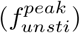 as a function of the noise amplitude when stimulating R-Precentral. (C) Average peak frequency of 81 unstimulated regions 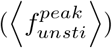 under different combinations of noise amplitude and stimulation sites. (D) (E) (F) Time series (upper panels) and power spectra (lower panels) of an unstimulated region (R-Precuneus) when stimulating the R-Lateral Orbitofrontal under different noise amplitudes (10^*−*7^, 10^*−*5^, 10^*−*3^). The blue and orange lines in the lower panels indicate the power spectrum before and during stimulation, respectively.

We then consider the peak frequency in unstimulated brain regions 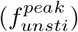. Unlike stimulated areas that directly receive external perturbations, these regions are indirectly affected through interactions with other regions. Fig. 3B shows the impact of the noise amplitude at a fixed stimulation site (R-Precentral). When *σ* is low, most regions exhibit high 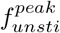, indicating that these regions effectively received the activity from the stimulated area. As *σ* increases, more regions exhibit low 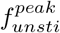 values, which implies that activity transmission is hindered. The behaviors of some other stimulation sites are provided in S3 Fig to validate the robustness of our results. Fig. 3C shows the average peak frequency of 81 unstimulated brain regions 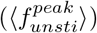 under different combinations of stimulation sites and noise amplitudes. The results show that *σ* induces a disturbance effect, reducing 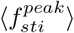 at all stimulation sites. Note that the effects of the noise amplitude differ at various stimulation sites. Some regions are more susceptible to noise and exhibit decreased 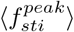 at relatively low *σ*. However, some other regions show the opposite behavior, indicating the complex interplay between the noise amplitude and network structure. Additionally, several large values at high *σ* can be observed in Fig. 3B, C because the system is already in an oscillatory state before stimulation, as shown in Fig. 2C.

To further elucidate the behaviors of unstimulated brain areas, we investigate the time series and power spectra of the R-Precuneus when stimulating the R-Lateral Orbitofrontal under different noise amplitudes as typical examples (Fig. 3D, E, F). We find that as the noise amplitude increases, the regional activity becomes increasingly irregular. Moreover, the power spectrum before stimulation increases and constrains that during stimulation as a lower bound. For robustness, we also show the behaviors of other unstimulated brain regions in S4 Fig.

We further explore the relationship between the noise amplitude and the network structure by investigating how oscillatory activity propagates from stimulated to unstimulated regions. In Fig. 4A, B, C, we show three typical examples of 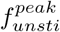 matrices under fixed noise amplitudes (*σ* = 10^*−*7^, 10^*−*5^, 10^*−*3^). Each element in the matrices represents the peak frequency of an unstimulated brain region (y-axis) under a specific stimulated area (x-axis). We observe that the 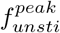 values heterogeneously decrease as *σ* increases. Thus, we are interested in which node pairs are more vulnerable to noise and how this behavior relates to structural properties. Fig. 4D presents the matrix of 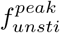 values averaged across noise amplitudes 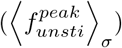. Intriguingly, this matrix is remarkably similar to the structural connectivity matrix, with a Spearman correlation coefficient of *r* = 0.93, *p <* 0.01 (Fig. 4E). Node pairs with strong direct connections tend to exhibit high 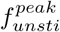 values in a large range of *σ*, indicating strong activity transmission capability. This result reveals that the antagonistic effects of the structural connection strength and noise amplitude shape activity propagation in terms of the peak frequency between stimulated and unstimulated areas to some extent. Note that this result also illustrates the rather small contribution of multistep paths due to the greater noise disturbance along the path. Oscillations before stimulation have little impact on this result (S5 Fig).

**Fig 4.**
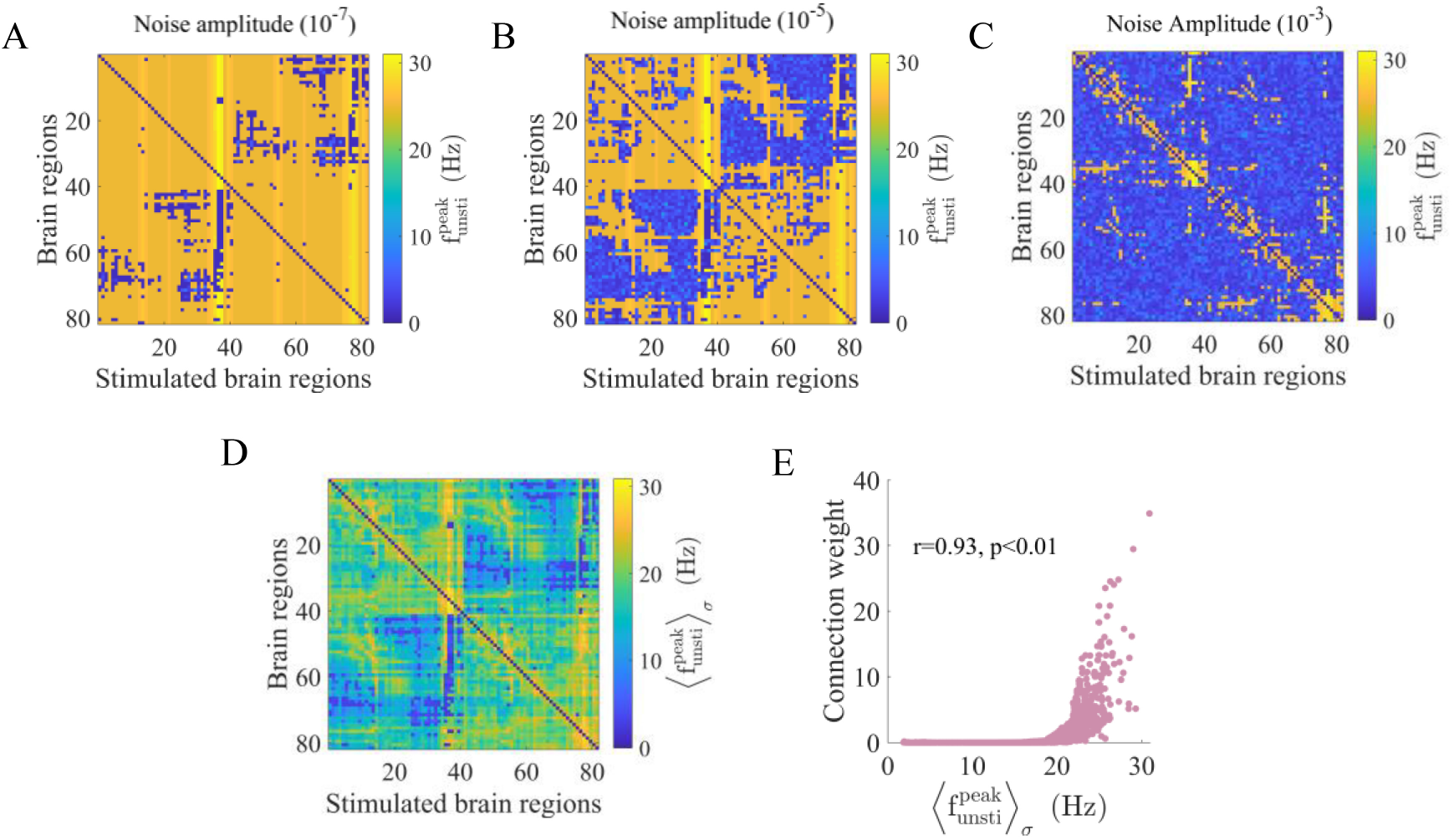
The high similarity between the peak frequency of unstimulated areas averaged across noise amplitudes and the structural connectivity. (A) (B) (C) The peak frequency 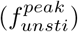 of unstimulated brain regions (y-axis) under various stimulated regions (x-axis) at different noise amplitudes (10^*−*7^, 10^*−*5^, 10^*−*3^). The diagonal elements are set to 0. (D) Peak frequency of unstimulated regions (y-axis) under different stimulation sites (x-axis) averaged across all noise amplitudes 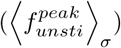, with the diagonal elements set to 0. (E) The positive Spearman correlation (*r* = 0.93, *p <* 0.01) between the matrix in (C) and the structural connectivity network.

### The heterogeneous impact of noise amplitude on structural degree alters network-level stimulation effects

In this section, we comprehensively investigate the network-level effects of stimulation (functional and structural effects) based on the functional connectivity matrix. Our goal is to explore how noise amplitude affects these network-level effects. Many previous studies have suggested that the structural degree of the stimulated region is an important feature for predicting and controlling stimulation effects [19, 37, 50]. How does noise amplitude influence the role of the structural degree? In particular, how does noise amplitude affect the relationship between the structural degree and functional or structural effects?

Fig. 5 shows the difference in functional networks before and during stimulation under three noise amplitudes (*σ* = 10^*−*8^, 10^*−*5^, 10^*−*2^) at three stimulation sites with various degrees (L-Pars Orbitalis, R-Precentral, and L-Caudate). We observe that as the noise amplitude increases, the changes in the functional networks decrease. When *σ* = 10^*−*8^, most node pairs in the networks exhibit large alternations. When *σ* = 10^*−*2^, the edge changes are small. Moreover, stimulating different regions leads to similar results in these two situations. When *σ* = 10^*−*5^, only some node pairs are influenced. The larger the degree of the stimulated region, the larger the range of alterations in the functional network. These results provide an intuitive illustration of how noise amplitude and structural degree collectively affect network-level effects. We also provide examples of brain stimulation in the oscillatory state, as shown in S6 Fig.

**Fig 5.**
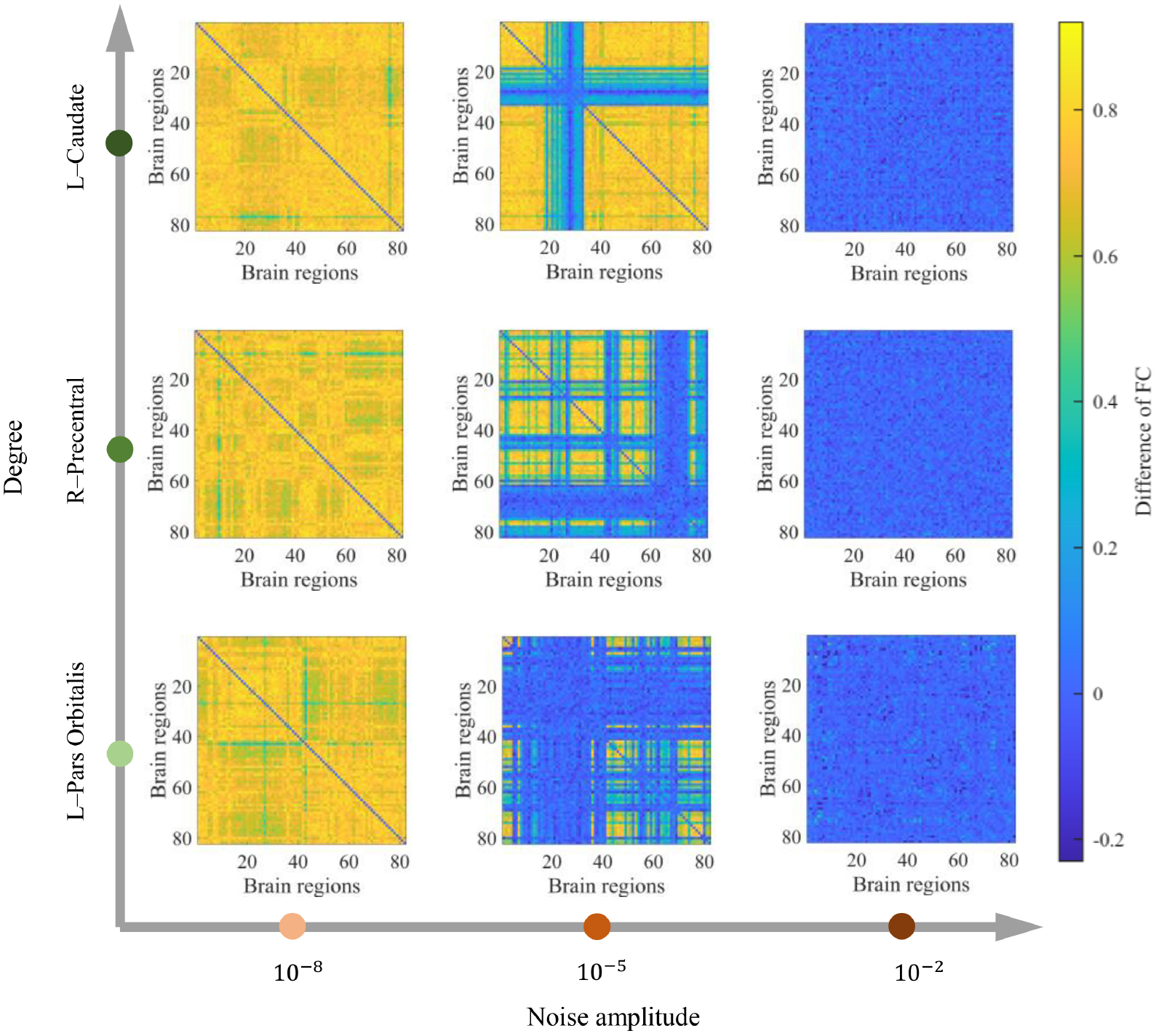
Examples of functional connectivity changes induced by stimulating different regions under various noise amplitudes. The brain regions L-Pars Orbitalis, R-Precentral, and L-Caudate with low, moderate and high structural degrees, respectively, are stimulated at low (10^*−*8^), moderate (10^*−*5^), and high (10^*−*2^) noise amplitudes. The matrices represent the differences in functional connectivity networks before and during local stimulation for one realization.

In Fig. 6, we use the functional effects to quantitatively investigate how functional brain networks are affected by external perturbations. We calculate the mean of the absolute values of the upper triangular elements in the functional connectivity difference matrices induced by local stimulation. Fig. 6A shows the functional effects under different combinations of noise amplitudes and stimulation sites. The impact of the noise amplitude can be separated into three distinct regimes. In the first regime (*σ <* 10^*−*8^), large functional effects are independent of the noise amplitude, and the local stimulation alters the system into a state that differs considerably from the prestimulation situation. In the second regime (10^*−*8^ *< σ <* 10^*−*3^), the functional effects gradually decrease as *σ* increases, showing disturbance effects. In the third regime (*σ >* 10^*−*3^), the functional effects are approximately 0, and the local stimulation has little impact on the brain. Moreover, there is obvious heterogeneity among stimulation sites under specific noise amplitudes, especially in the second regime, indicating the effect of brain structure. Additionally, note that the stimulation sites exhibit different levels of resistance to noise. To understand the role of the structural degree and its interaction with the noise amplitude, we exhibit the functional effects versus the noise amplitude separately for all stimulation sites and color the effects according to the corresponding degree, as shown in Fig. 6B. We found that the larger the degree of the stimulation site, the larger the functional effects under a fixed noise amplitude and the stronger the noise amplitude required to reduce functional effects. This result demonstrates the heterogeneous effect of noise on the degree, i.e., regions with large degrees not only have a high capacity to influence brain dynamics but also show strong resistance to noise.

**Fig 6.**
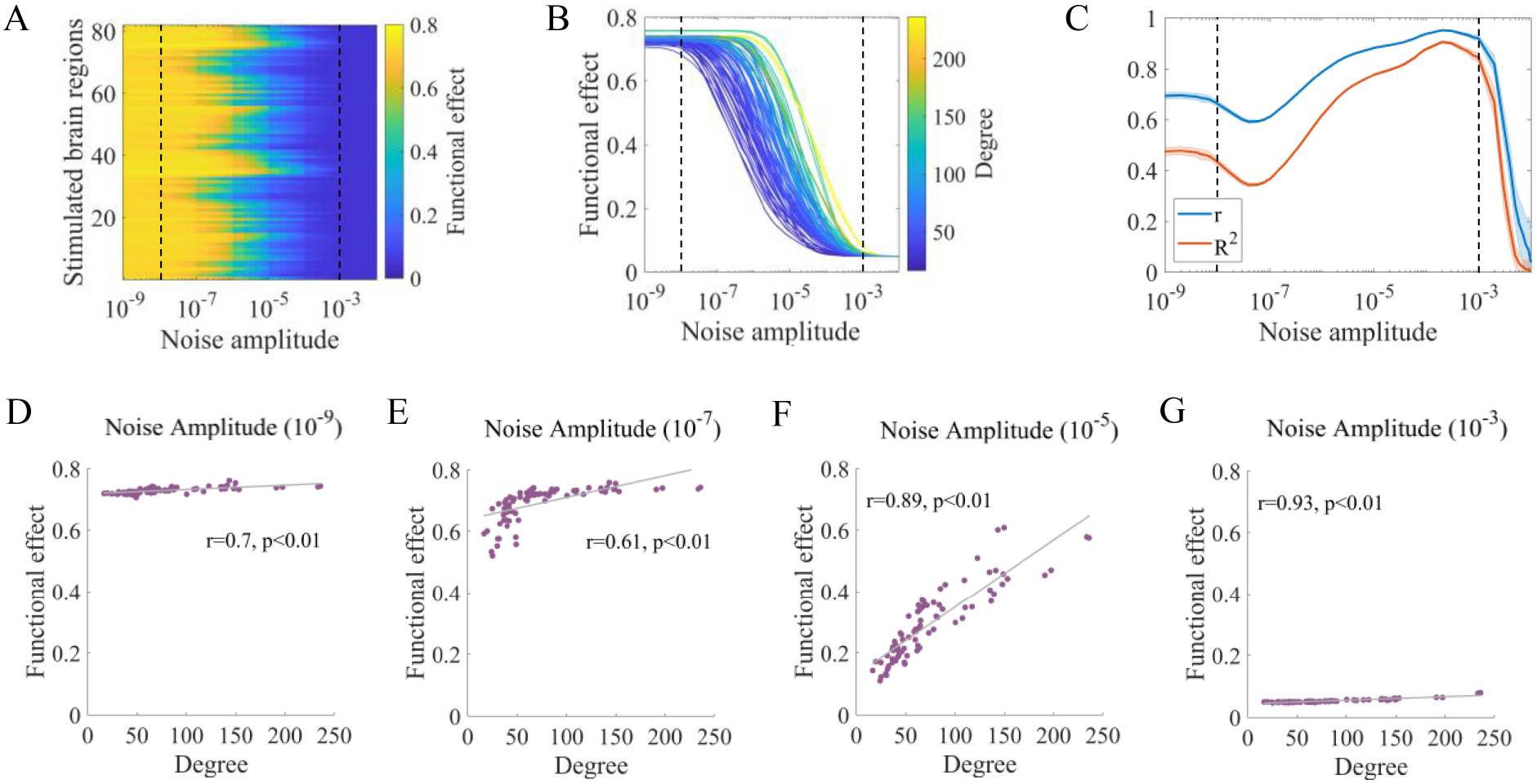
Noise amplitude and structural degree jointly affect the functional effects of stimulation. (A) Functional effects under different combinations of stimulation sites and noise amplitudes. (B) Functional effects as a function of noise amplitude for all stimulation sites, ranked by structural degree. Note that the values in (A) and (B) represent the ensemble averages of 30 realizations of the corresponding measures. (C) Pearson correlation coefficient (*r*) and adjusted coefficient of determination (*R*^2^) between functional effects and structural degree as a function of noise amplitude. The solid lines and shaded areas describe the ensemble averages and standard deviations of 30 realizations of the corresponding measures. (D) (E) (F) (G) Snapshots of the functional effects under different noise amplitudes for one realization. (D) Noise amplitude = 10^*−*9^, Pearson’s *r* = 0.7, *p <* 0.01. (E) Noise amplitude = 10^*−*7^, Pearson’s *r* = 0.61, *p <* 0.01. (F) Noise amplitude = 10^*−*5^, Pearson’s *r* = 0.89, *p <* 0.01. (G) Noise amplitude = 10^*−*3^, Pearson’s *r* = 0.93, *p <* 0.01. The gray lines represent the linear fits of data points estimated by ordinary least squares.

In Fig. 6C, the Pearson correlation coefficient *r* between the functional effects and structural degree as well as the adjusted coefficient of determination *R*^2^ estimated via an ordinary least squares method are presented as functions of the noise amplitude. High absolute values of *r* and *R*^2^ indicate that the functional effects are highly linearly correlated with and well fitted by the corresponding degree. Initially, *r* shows intermediate values and remains approximately constant (first regime); then, *r* decreases to a local minimum and increases to a global maximum (second regime) and finally rapidly declines to 0 (third regime). *R*^2^ shows a similar trend. We also provide typical snapshots of the functional effects under different noise amplitudes in Fig. 6D-G (*σ* = 10^*−*9^, 10^*−*7^, 10^*−*5^, 10^*−*3^), and snapshots under other noise amplitudes are shown in S7 Fig. These results show that noise amplitude nonmonotonically modulates the relationship between functional effects and structural degree. When *σ* = 10^*−*5^, we observe a high positive association for functional effects, which is consistent with a previous study [37]. However, only a moderate level of correlation was observed when the noise had little impact on brain dynamics (Fig. 6D), highlighting its nontrivial role. Furthermore, increasing the noise amplitude in the second regime could progressively enhance the correlation and the predictability of functional effects.

To better understand these behaviors, we provide further explanations from the perspective of the underlying dynamical mechanisms. In general, functional effects depend on the interplay between network structure and noise amplitude. Under the small noise amplitude which has little impact on neural activity (Fig. 6D), the intrinsic network structure plays a major role. Stimulation sites with larger degrees tend to have more neighbors with higher connection weights and shorter transmission delays than regions with smaller degrees [51], thereby facilitating more effective information transmission. Therefore, although the functional effects are all quite high, they are moderately correlated with the structural degree. Following previous analyses, stimulation sites with small degrees limit information transmission, causing the propagation of downstream activity sensitive to the increased noise amplitude. When the noise amplitude is slightly larger (Fig. 6E), the functional effects induced by stimulation to small-degree regions are reduced, while the functional effects are essentially unchanged for regions with large degrees. This relationship becomes nonlinear and the linearity diminishes. As the noise amplitude gradually increases (Fig. 6F, G), the downstream activity transmission is significantly hindered. However, neighboring areas are less affected due to their direct connections with stimulated regions. This analysis indicates that the structural degree plays a progressively important role in predicting the response to stimulation, leading to the linearity increasing, although the functional effects decrease.

An important feature of the brain is that a relatively static network structure supports complex dynamic functions. Therefore, the structure-function coupling can reflect the network-level state. In Fig. 7, we study the alterations in the extent to which brain function is limited by the network structure through structural effects. We calculate the difference in the Pearson correlation coefficient between the structural and functional connectivity matrices before and during stimulation. Fig. 7A exhibits the structural effects as a function of the noise amplitude and stimulation site. We evaluate the structural effects according to the three regimes shown in Fig. 6. In the first regime, the structural effects show moderate values and are independent of the noise amplitude. Local stimulation causes a temperate increase in the similarity between the structural and functional connectivity. In the second regime, the structural effects of each region increase to their peak values under large noise amplitudes, indicating that function connectivity is more constrained by the network structure. In the third regime, all structural effects decrease to small values near 0. Moreover, we observe heterogeneous behaviors among stimulation sites, especially in the second regime, indicating the crucial role of the interaction between noise amplitude and network structure. Following previous analyses, we present the structural effects as a function of the noise amplitude separately for all stimulation sites and color the results according to the corresponding degree, as shown in Fig. 7B. We find that the structural degree is not only related to the structural effects under a fixed noise amplitude but also positively correlated with the noise amplitude required to achieve the peak values in the second regime. This result indicates that the noise amplitude has a diverse influence on the degree, leading to the different performance of the structural constraints across the stimulated regions.

**Fig 7.**
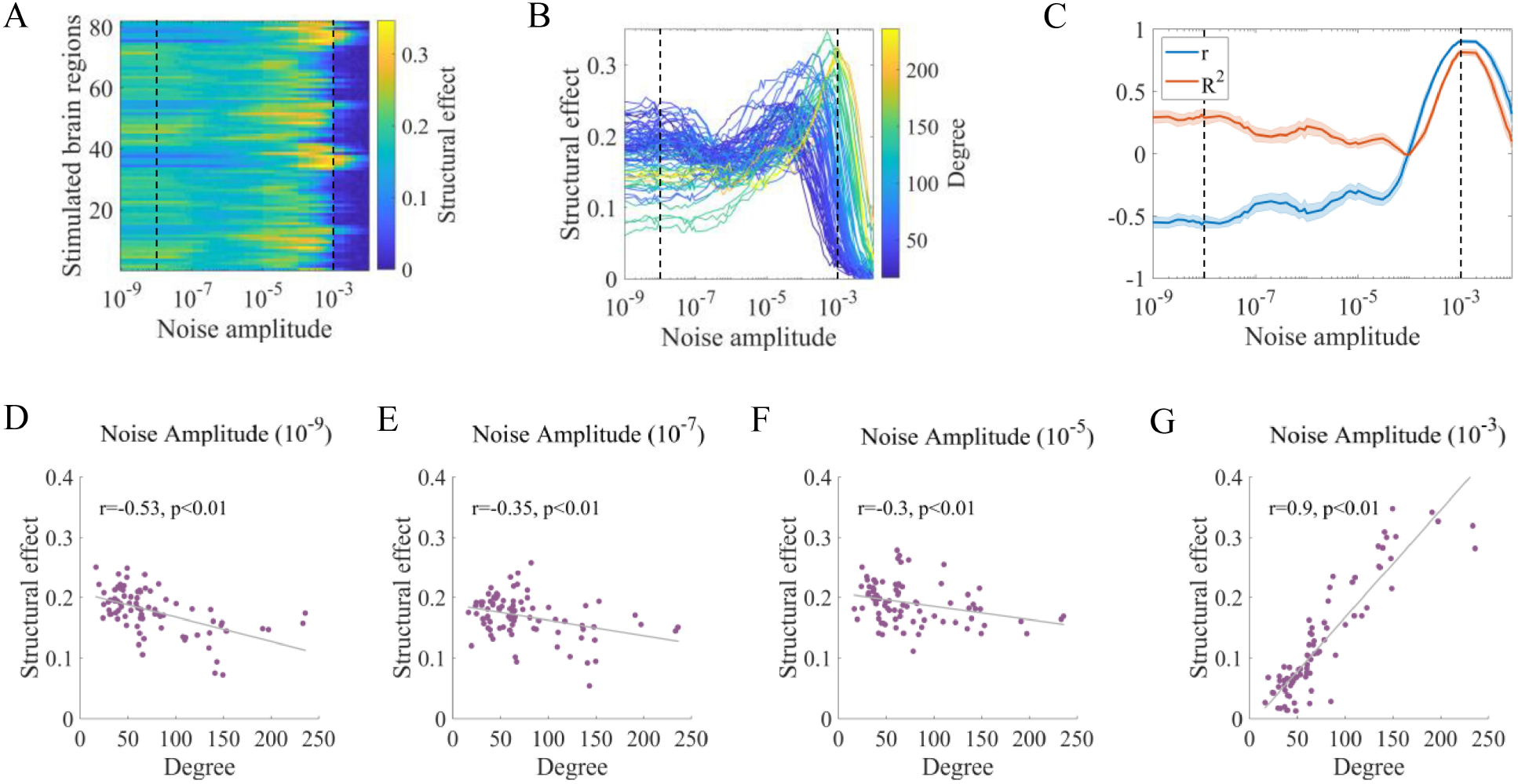
Noise amplitude and structural degree jointly influence the structural effects of stimulation. (A) Structural effects under different combinations of stimulation sites and noise amplitudes. (B) Structural effects as a function of noise amplitude for all stimulation sites, ranked by structural degree. Note that the values in (A) and (B) represent the ensemble averages of 30 realizations of corresponding measures. (C) Pearson correlation coefficient (*r*) and adjusted coefficient of determination (*R*^2^) between structural effects and structural degree as a function of the noise amplitude. The solid lines and shaded areas describe the ensemble averages and standard deviations of 30 realizations of corresponding measures. (D) (E) (F) (G) Snapshots of structural effects under different noise amplitudes for one realization. (D) Noise amplitude = 10^*−*9^, Pearson’s *r* = −0.53, *p <* 0.01. (E) Noise amplitude = 10^*−*7^, Pearson’s *r* = −0.35, *p <* 0.01. (F) Noise amplitude = 10^*−*5^, Pearson’s *r* = −0.3, *p <* 0.01. (G) Noise amplitude = 10^*−*3^, Pearson’s *r* = 0.9, *p <* 0.01. The gray lines represent the linear fits of data points estimated by ordinary least squares.

Analogous to Fig. 6C, we show the Pearson correlation coefficient *r* and the adjusted coefficient of determination *R*^2^ for the structural effects in Fig. 7C. In the first regime, *r* and *R*^2^ exhibit moderate values with opposite signs. In the second regime, *r* remains negative and then rapidly increases to a positive value near 1. *R*^2^ shows similar behavior but with positive values. In the third regime, these metrics decrease rapidly to low values. Typical snapshots of the structural effects under different noise amplitudes are shown in Fig. 7D-G (*σ* = 10^*−*9^, 10^*−*7^, 10^*−*5^, 10^*−*3^). Snapshots under other noise amplitudes are illustrated in S8 Fig. These results show that the relationship between the structural effects and structural degree is nonmonotonically affected by the noise amplitude. When *σ* = 10^*−*5^, we find a weak negative association for the structural effects, which is consistent with a previous study showing poor predictability [37]. Nevertheless, our results indicate that when the noise amplitude is larger, the structural effects are highly correlated with and well predicted by the structural degree. Specifically, increasing the noise amplitude in the second regime can enhance the structural effect correlations and even change its sign from negative to positive.

To better understand these nonmonotonic changes, we present interpretations based on fundamental dynamical mechanisms. The structural effects reflect the similarity between structural and functional connectivity, which is modulated by the stimulation site and noise amplitude. Under relatively low noise amplitudes (Fig. 7D-F), stimulating regions with large degrees produces high functional connectivity in most node pairs, indicating the low correspondence between the structural and functional connectivity and the reduced structural effects. In contrast, stimulating regions with small degrees leads to more node pairs showing low functional connectivity. As the structural connection weights reflect the information transmission ability to some extent, the low functional connectivity is more likely to be found at node pairs with low connection weights, therefore inducing higher structural constraints. Consequently, the structural effects are negatively correlated with the structural degree. As the noise amplitude increases (Fig. 7G), disturbance effects are enhanced. Most functional connectivity shows values near 0 when stimulating small-degree areas, indicating the low correspondence between the structural and functional connectivity and reduced structural effects. For large-degree stimulation sites, more functional connectivity shows high values, which is more likely to be found at node pairs with high structural connection weights, resulting in high structural constraints. Hence, the structural effects are positively correlated with the structural degree.

### Behaviors of cognitive systems in the structure–function landscape are jointly modulated by noise amplitude and the average system degree

In this section, we categorize brain regions into cognitive systems, stimulate areas in single systems, and study system behaviors based on local stimulation effects. We employ a coarse-grained classification with four cognitive systems: the sensory and association (SA) system, higher-order cognitive (HOC) system, medial default mode (MDM) system, and subcortical system [52]. The stimulation effects are evaluated according to the functional and structural effects. Previous research has shown that the cognitive functions of brain systems are related to their stimulation effects [37, 53]. Here, we focus on the following questions: How does noise amplitude influence the stimulation effects of different cognitive systems? What is the association between noise-induced impacts and the system-level network structure?

In Fig. 8A, B, we present the mean and standard deviation of the functional effects induced by stimulating regions in single cognitive systems (⟨*fe*⟩ and *std* (*fe*)) as a function of the noise amplitude. Fig. 8A shows that ⟨*fe*⟩ of each system nonlinearly decreases as the noise amplitude increases. The subcortical system shows the highest noise amplitude required to reduce ⟨*fe*⟩, followed by the MDM, HOC, and SA systems. Moreover, the ⟨*fe*⟩ values of different cognitive systems follow the same order at various noise amplitudes, indicating a relatively consistent pattern in terms of the impacts on the functional configuration. According to Fig. 8B, as the noise amplitude increases, *std* (*fe*) first increases to a global maximum and then decreases to 0. The SA system is the first to reach its peak value, followed by the HOC and MDM systems, and finally, the subcortical system, indicating the different levels of flexibility of functional effects in distinct systems at various noise amplitudes.

**Fig 8.**
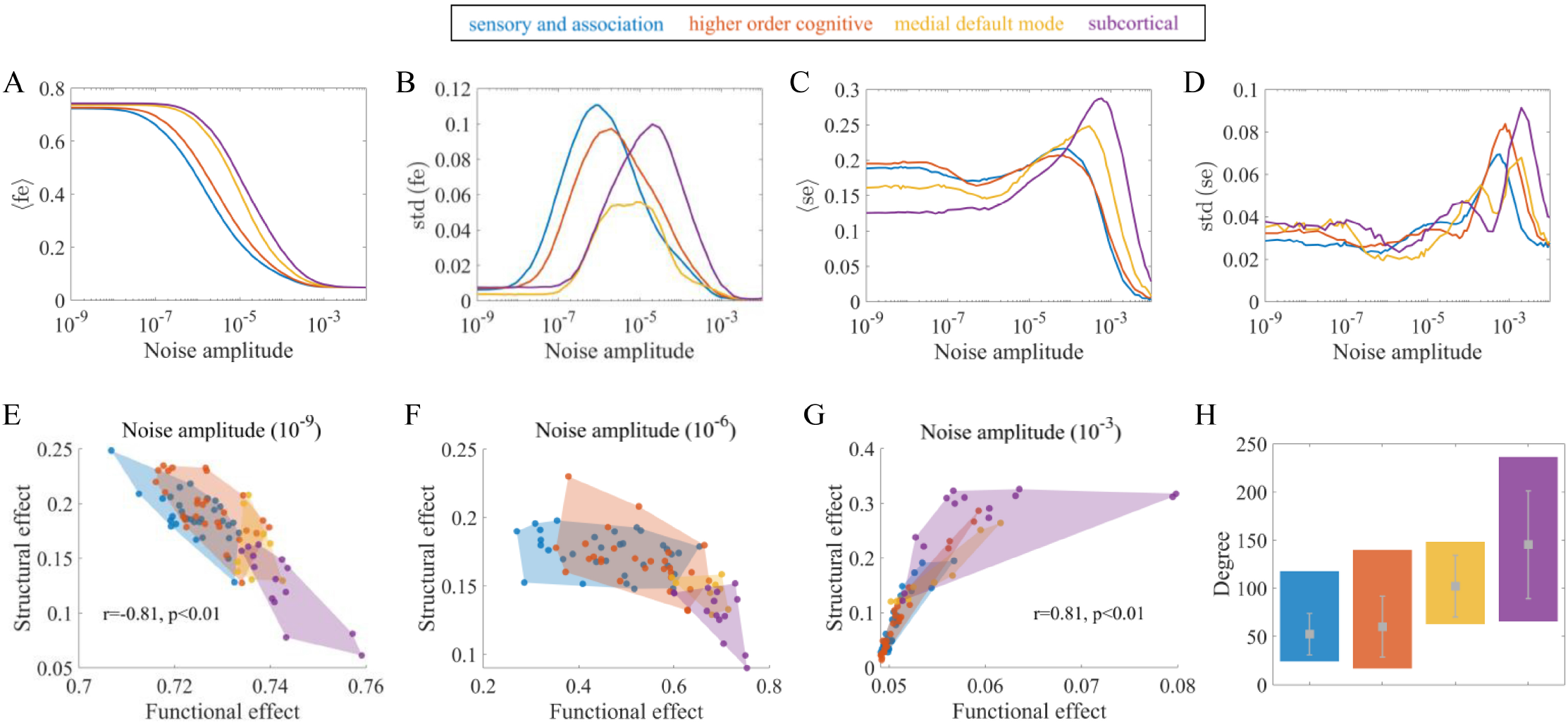
The associations between behaviors of cognitive systems and noise amplitude and the average system degree. (A) Mean of the functional effects induced by stimulating regions in a single system (⟨*fe*⟩) versus the noise amplitude. (B) Standard deviation of the functional effects induced by stimulating regions in a single system (*std* (*fe*)) versus the noise amplitude. (C) Mean of the structural effects induced by stimulating regions in a single system (⟨*se*⟩) versus the noise amplitude. (D) Standard deviation of the structural effects induced by stimulating regions in a single system (*std* (*se*)) versus the noise amplitude. (E) (F) (G) Structural effects versus functional effects of cognitive systems at various noise amplitudes (10^*−*9^, 10^*−*6^, 10^*−*3^). Note that the brain regions are grouped into 4 cognitive systems, as indicated by the different colors. The colored areas represent the convex hulls of data points in the systems. The lines and points reflect the measures averaged over 30 realizations. (H) Structural properties of each cognitive system. The colored bars indicate the maximum and minimum structural degree of regions in the systems. The gray dots and error bars represent the mean and standard deviation of the structural degree in the systems.

Fig. 8C, D shows the mean and standard deviation of the structural effects in each cognitive system (⟨*se*⟩ and *std* (*se*)). Fig. 8C presents that as the noise amplitude increases, ⟨*se*⟩ initially shows a moderate value, then increases to a maximum and finally decreases. The noise amplitude corresponding to the peak value of ⟨*se*⟩ in the subcortical system is larger than that in the SA and HOC systems, while the MDM system shows a moderate value. The ⟨*se*⟩ values of the different systems follow the same order under large noise amplitudes. Nevertheless, the situation shows the opposite trend under small noise amplitudes, with the SA and HOC systems showing the largest ⟨*se*⟩ values, indicating the altered profiles of cognitive systems at various noise amplitudes in terms of structural constraints. The *std* (*se*) also shows nonmonotonic trends with global peaks, as presented in Fig. 8D. The SA and HOC systems reach their global maximums at lower noise amplitudes than the MDM and subcortical systems, leading to different expressions of the variability of structural constraints across noise amplitudes.

Fig. 8E-G presents the locations of cognitive systems in the structure–function landscape under different noise amplitudes (*σ* = 10^*−*9^, 10^*−*6^, 10^*−*3^). According to Fig. 8E, the functional effects are negatively related to the structural effects. In contrast to the other systems, regions in the SA and HOC systems have smaller impacts on functional configurations and are more constrained by the structural network, while stimulating subcortical areas shows the opposite behavior. Fig. 8F shows the nonlinear relationship between functional and structural effects, which is comparable to the inverted U-shaped curve discussed in previous research [37]. Stimulating regions in the SA and HOC systems results in high variability of functional effects which are highly constrained by the structural network. In contrast, regions in the MDM and subcortical systems exhibit large functional effects and small structural effects. Fig. 8G shows a positive association between these two measures. Stimulating subcortical areas are more structurally constrained than the SA and HOC systems. Overall, these results indicate that the noise amplitude not only alters the stimulation effects of different cognitive systems but also their relations with each other. The locations of different cognitive systems under other noise amplitudes are shown in S9 Fig.

Finally, we present the properties of the structural degree of each cognitive system in Fig. 8H. We observe that the subcortical system has the highest average degree, followed by the MDM, HOC and SA systems. This order is consistent with many cognitive system ranks of noise-related stimulation effects, such as values and resistance to noise in Fig. 8A, the values in Fig. 8C, and the noise amplitude corresponding to the peak values in Fig. 8B-D. These results highlight the mechanism of the mean structural degree as an intrinsic property of cognitive systems in modulating noise-induced effects and indicates that the normal function of cognitive systems is jointly dependent on the noise amplitude and network structure.

## Discussion

Understanding the effects of local stimulation is essential for revealing the causal relationship between neural activity and cognition and promoting clinical applications for regulating or restoring brain function [10–16]. Although many efforts have been made in exploring fundamental principles, the global response to stimulation is not fully understood. Given the evidence that noise contributes to the variability across subjects and that the modulation of neural activity induced by noninvasive stimulation may alter the signal-noise relationship [35, 36], we hypothesize that noise amplitude is a crucial factor affecting neural activity patterns during stimulation.

Inspired by the past theoretical work [27, 37, 38, 40], we simulated a whole-brain biophysical model under different combinations of noise amplitudes and stimulation sites to elucidate the associations between noise amplitude and the impacts of local stimulation and, more importantly, the interplay between noise amplitude and network structure. We first determined an optimal value for the global coupling strength before stimulation and then assessed the effects of regional perturbations. From a regional perspective, local perturbations increased the peak frequency of neural activity, similar to previous findings in natural and experimental stimulation studies [36, 54]. We observed that noise amplitude has little impact on the peak frequency of stimulated brain regions but reduced that of unstimulated areas. In addition, we found a high similarity between the peak frequency matrix for unstimulated areas averaged across various noise amplitudes and the structural network. From a network perspective, we quantified the effects of stimulation by examining the overall changes in functional connectivity (functional effects) and the variations in structure–function coupling (structural effects). We observed that noise amplitude nonlinearly decreased functional effects and nonmonotonically modulated structural effects. Crucially, we found that noise amplitude nonmonotonically altered both the Pearson correlation coefficient and adjusted coefficient of determination between the structural degree and functional and structural effects, which has potential utility in better predicting and controlling therapeutic performance. Finally, we showed that the behaviors of different cognitive systems in the landscape of functional and structural effects depended on the interplay between the noise amplitude and system-level average structural degree.

We first provide a discussion on the impacts of noise amplitude. Noise is inevitable and common in the brain and shows both detrimental effects and potential benefits [28, 55]. We showed that under a small noise amplitude, the neural activity patterns altered by perturbations easily spread throughout the network, resulting in high peak frequencies in most brain areas and strong functional couplings as indicated by functional effects. This behavior is an abnormal manifestation resembling the state induced by generalized epilepsy, which is often associated with enhanced interregional synchronization and is not conducive to effective information processing [56, 57]. In contrast, large noise amplitudes disrupted the transmission of neural activity; thus, most peak frequencies and functional couplings could not be enhanced by local stimulation. Previous research has shown that many brain disorders such as autism, schizophrenia and cognitive dysfunction induced by aging or fibromyalgia are linked to increased neural noise [58–62]. These decreased signal-to-noise ratios have been shown to contribute to power spectrum density changes, decreased oscillatory coherence, and network communication errors [63], which is similar to our findings. Furthermore, stimulation with a moderate noise amplitude tends to elicit temperate effects, influencing only part of the brain, which is consistent with a previous study, emphasizing the importance of the partial synchronization state in cognition [53]. These findings indicate that neural noise amplitude may be crucial in affecting widespread changes in regional activity and functional interactions caused by stimulation. Our results augment the literature on how noise affects neural communication dynamics from the perspective of local stimulation and support the notion that an appropriate noise amplitude is essential for maintaining brain functions such as receiving external environmental stimuli, performing internal information processing and executing normal cognitive functions [29–34, 64, 65].

Our analyses also demonstrated that noise amplitude nonmonotonically modulates the dependence of brain function on structure, i.e., structural effects. This finding is reminiscent of a recent study showing that changes in neural noise in some brain regions drive structure–function decoupling [66]. Intriguingly, increasing the noise amplitude in a specific range of the second regime improved the similarity between structural and functional connectivity, which reflects the complex behaviors associated with the structure–function relationship and may be relevant to normative brain dynamics [67].

Recent studies have suggested that ketamine anesthesia increases the randomness of neural activity, likely associated with a decreased neural signal-to-noise ratio [68, 69] and has variable effects across brain regions [70, 71], which leads to different stimulation impacts. For example, stimulation to the ventral tegmental area under ketamine anesthesia elicits smaller network activation than in the awake state [72]. In contrast, stimulation to the parietal cortex shows similar distal effects in both states [73]. Analogously, in this work, we found a heterogeneous effect of noise amplitude on stimulation sites in terms of both regional and network-level stimulation effects. Here, we conceptualize stimulation sites as structural network properties and then discuss the interaction between the noise amplitude and network structure.

From a regional perspective, the positive correlation between the peak frequency of unstimulated areas under different stimulation sites averaged across noise amplitudes and the structural connectivity implies an antagonistic effect between structural connection strength and noise amplitude. The gradually increasing noise amplitude acts as a high-pass filter on the structural brain network and is more likely to impede communication between node pairs with weak weights. The relationship between brain structure and function is one of the most important challenges in neuroscience [74]. Many studies have focused on predicting brain function according to network structure [75–77]. In this work, by leveraging noise amplitude and local perturbations, we derive information about the structural network according to the functional data obtained from numerical simulations, which improves our understanding of structure–function associations in the brain.

It is of great interest to predict and control network-level responses to stimulation [10, 78], and many studies have proposed the structural degree as an important property [19, 50]. We observed that under a moderate noise amplitude (10^*−*5^), the structural degree showed a strong positive correlation with functional effects and a weak negative correlation with structural effects. This result is in accordance with a previous study showing that the structural degree mainly controls functional effects and that structural effects could not be easily predicted based on whether an area was a hub or nonhub [37]. Furthermore, we found that the noise amplitude modulates the association between the structural degree with functional and structural effects in a nontrivial way. In particular, there was only a moderate level of correlation when the noise amplitude had little impact on brain dynamics. Thus, the fact that the structural degree could serve as a good predictor of functional effects is not only an intrinsic property of the network structure but also attributed to the noise amplitude. This result deepens our understanding of the structural degree and emphasizes the significance of considering specific dynamical processes when investigating the role of structure [79]. Our result also indicates that increasing the noise amplitude within specific ranges of the second regime improves the association for functional and structural effects and improves predictability and controllability, which shows important potential applications in research experiments and disease treatments related to stimulation. The growing noise amplitude even influences the sign of the correlation for structural effects, changing it from negative to positive. These analyses highlight the necessity of considering the coupling of multiple factors, including the brain structure and noise amplitude, to comprehensively understand the global effects of stimulation. A recent study also supports this notion by showing that both stimulation sites and brain collective oscillatory states can affect the widespread impacts of focal stimulation [27].

Previous research has shown that brain regions exhibit specific trade-offs between functional and structural effects that are linked to their cognitive function [80, 81]. As a result, cognitive systems, which are defined as subgraphs of the brain, occupy various locations in the structure–function landscape [37, 53]. At a moderate noise amplitude (10^*−*6^), we observed that the nonlinear associations between the functional and structural effects of different systems were consistent with the findings reported in [37]. Moreover, our results showed that noise amplitude reduced the average functional effect and nonmonotonically modulated the average structural effect, as well as the variability of these two effects, therefore altering the positions of the systems in the structure–function landscape, potentially implying a functional disorder. Previous research has found abnormalities in functional connectivity and its variability in the default mode and somatomotor networks in diseases characterized by aberrant neural noise (such as schizophrenia and autism) [82–84], which may be associated with motor and cognitive dysfunction. Furthermore, we observed that cognitive systems express specific organizing principles at various noise amplitudes. Taking the average functional effect as an example, the subcortical system showed the highest functional effects and the strongest resistance to noise, while the sensory and association system exhibited the opposite trend. This result may be explained by the fact that the subcortical system tends to play a global role in network dynamics, facilitating communication between brain areas, whereas the sensory and association system tends to play a specialized role, working in segregation and activating only a small part of the brain [85, 86]. These heterogeneous behaviors at various noise amplitudes were largely attributed to the system-level mean structural degree, which again emphasizes the important role of both noise amplitude and network structure in shaping brain dynamics during stimulation.

We next discuss several potential clinical applications about the combined effects of noise amplitude and network structure. Due to the individual differences in noise amplitude, our results contribute to understanding the highly variable consequences of stimulation and facilitating the development of personalized healthcare approaches [25, 26]. In particular, for patients with brain disorders characterized by abnormal neural noise such as autism and schizophrenia [58–60], therapists need to carefully consider the role of noise amplitude when treating with local stimulation. In addition, appropriately adjusting the noise amplitude could increase the correlations between the structural degree and network effects, which supports the use of linear control theory and may improve outcome predictions [87, 88]. Note that there is a trade-off between functional effects and their predictability. The maximum *r* and *R*^2^ values are achieved at the expense of functional effects, and stimulation induces only a small fraction of changes in functional networks, regardless of the stimulation site. In contrast, at lower noise amplitudes, local perturbations produce broad changes in functional connectivity. However, the *r* and *R*^2^ values are reduced to some extent. This result suggests that researchers should carefully tune the noise amplitude according to practical needs to balance the range of impact and the predictability.

Finally, we provide several limitations of the present study and prospects for future work. Following previous studies [27], we used a structural brain network consisting of 82 areas based on a low-resolution atlas. Although the relatively small number of nodes and connections is beneficial for computationally dense simulations of key variables, this approach may ignore important structural information at finer scales. Moreover, the group-representative connectome precludes the exploration of network differences across individuals. In addition, the main goal of this work is to demonstrate from a general perspective how noise amplitude influences the effects of local stimulation. We chose the canonical Wilson-Cowan neural mass for brain dynamics, including a constant excitation as regional perturbations, and configured the global coupling strength such that the neural activity lies just before the high-activity oscillatory state, which is assumed to support empirical brain functions and provides maximal flexibility to perturbations [38, 47, 48]. However, this computational model is a simplification of the empirical situation and thus cannot perfectly describe the patterns of neural activity [89]. Future work could consider more realistic improvements, such as incorporating complex stimulation protocols [90], additional regional heterogeneity [91–93], and synaptic plasticity [94]. Importantly, these results should be tested experimentally using local stimulation under different noise levels to ensure the validity of the biological insights provided by the model. For example, future research may cautiously use psychedelics such as ketamine and LSD to enhance entropy in brain areas, thereby leading to more disordered states, which are associated with changes in neural noise [95]. Moreover, our study could be viewed as an extension of the state-dependent stimulation, with noise amplitude reflecting brain states. In the future, the intrinsic activity of other states, such as sleep and working memory, can be considered to investigate how these states affect the stimulation outcomes.

## Materials and methods

### Empirical data

We utilize a group-level anatomical brain network mentioned in a previous study [27], which is derived by implementing deterministic tractography algorithms for diffusion-weighted MRI of 30 healthy subjects [96, 97]. This weighted and undirected structural connectome contains 68 cortical areas and 14 subcortical areas, which are defined according to a relatively coarse-grained atlas [42]. The connection weights are calculated as the number of white matter streamlines between regions and are normalized by the geometric mean of their volumes. The group-representative distance matrix, in which elements represent the Euclidean distance between the centers of brain regions averaged over all subjects, is obtained from the same dataset. More information about the participant demographics and the acquisition and processing procedures of neuroimaging data is presented in [27].

### Network model of brain dynamics

To simulate brain activity, we employ a nonlinear neural mass model, which has been widely used to investigate brain functions [98, 99]. Each brain region is composed of both excitatory and inhibitory neural populations and is governed by the Wilson-Cowan dynamics [43]. Brain areas are coupled through the structural connectivity described above, with distance-dependent time delays. In accordance with previous works [27, 37, 44], anatomical connections link only excitatory populations in different brain areas.

The activity of the *i*^*th*^ brain region is controlled by the following equations:

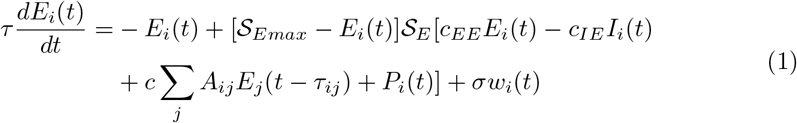

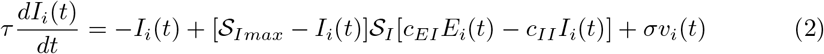

where *E*_*i*_(*t*) and *I*_*i*_(*t*) represent the mean firing rate of excitatory and inhibitory pools in brain region *i*, and *τ* is a time constant for both populations. The sigmoidal transfer functions 𝒮_*E*_ and 𝒮_*I*_ of the excitatory and inhibitory populations are described by

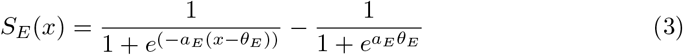

and

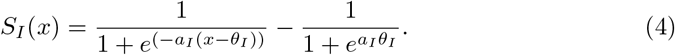

The fixed parameters *a*_*E*_ and *a*_*I*_ determine the maximal values of the slope, while *θ*_*E*_ and *θ*_*I*_ represent the positions of the maximum slope of the activation functions for each pool [43].

In each brain region, the excitatory pool receives local excitation from itself with strength *c*_*EE*_ and local inhibition from the inhibitory pool in the same region with strength *c*_*IE*_, as well as long-range excitation from excitatory pools in other regions through anatomical connections *A*_*ij*_ with global coupling strength *c* and external input *P*_*i*_(*t*). Due to the large distance between brain areas and the limited transmission speed, we also consider the time delay between regions *i* and *j* as *τ*_*ij*_, which is given by 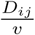. *D*_*ij*_ is an element in the group-representative distance matrix *D*, which indicates the mean Euclidean distance between regions *i* and *j*, and *v* is the velocity of signal conduction. The inhibitory pool receives only local excitation from the excitatory pool in the same region with strength *c*_*EI*_ and local inhibition from itself with strength *c*_*II*_. In addition, the excitatory and inhibitory populations are disturbed by random noise *w*_*i*_(*t*) and *v*_*i*_(*t*) with amplitude *σ. w*_*i*_(*t*) and *v*_*i*_(*t*) both follow standard Gaussian distributions.

The values of the model parameters used in this study are shown in Table 1 and are consistent with those used in previous research [37]. The local perturbation is set as a persistent excitation with intensity *P*_*i*_ = 1.25 for stimulated region *i* and an intensity of 0 for other areas. For an isolated brain area with the parameters shown in Table 1, external stimulation with an intensity of 1.25 causes a transition from a fixed point to the limit cycle regime [27, 37, 53]. The frequency of rhythmic activity for the stimulated region is approximately 20 Hz, which is essential in oscillatory neuronal dynamics [45, 46]. By tuning the strength of the excitatory input from other regions in the brain network, the global coupling strength *c* affects the system state, as reflected in the sudden increase in the mean firing rate in most regions, indicating the dynamic transition from a low-activity steady state to a high-amplitude oscillatory state. Note that noise amplitude *σ* is the parameter of interest. Therefore, we consider the global coupling strength *c* ∈ [0.01, 0.3] in steps of 0.005 and the noise amplitude *σ* ∈ [10^*−*9^, 10^*−*2^] in a log manner such as 10^*−*9^, 2 × 10^*−*9^, …, 9 × 10^*−*9^, 10^*−*8^, 2 × 10^*−*8^, … .

**Table 1.** Values of model parameters.

| Parameter | Description | Value |
| --- | --- | --- |
| $\tau$ | Time constant | 8 ms |
| $c_{EE}$ | Local excitatory-to-excitatory coupling strength | 16 |
| $c_{IE}$ | Local inhibitory-to-excitatory coupling strength | 12 |
| $c_{EI}$ | Local excitatory-to-inhibitory coupling strength | 15 |
| $c_{II}$ | Local inhibitory-to-inhibitory coupling strength | 3 |
| $a_E$ | Proportional to the excitatory maximum slope | 1.3 |
| $a_I$ | Proportional to the inhibitory maximum slope | 2 |
| $\theta_E$ | Position of the excitatory maximum slope | 4 |
| $\theta_I$ | Position of the inhibitory maximum slope | 3.7 |
| $\mathcal{S}_{E_{max}}$ | Maximum of the excitatory activity function | 0.9945 |
| $\mathcal{S}_{I_{max}}$ | Maximum of the inhibitory activity function | 0.9994 |
| $c$ | Global coupling strength | 0.01-0.3 |
| $v$ | Velocity of signal conduction | 10 m/s |
| $P_i$ | External stimulation intensity | 1.25 |
| $\sigma$ | Noise amplitude | $10^{-9} - 10^{-2}$ |

### Simulation details

Due to the large range of *σ*, we integrate the set of stochastic differential equations described above using the Euler-Maruyama scheme with a sufficiently small step (*dt* = 5 ×10^*−*6^*s*). We select a constant initial condition for all regions following previous work [37, 53]. The simulations are first performed for 2 seconds without stimulation under different global coupling strengths and noise amplitudes to determine the appropriate *c* (Fig. 2). Then, we rerun the simulations for 3 seconds under different noise amplitudes and stimulation sites, with the local perturbation applied for 2-3 s and a fixed global coupling strength. We perform each simulation 30 times and discard the first second of neural activity due to the initial instability. We mainly focus on the excitatory firing rate *E*_*i*_ (*t*) in each region [37, 44, 98, 99] and downsample these time series to a resolution of 1×10^*−*3^ s.

### Analyses metrics

The brain shows remarkably different dynamic performance in the before-(1-2 s) and during-stimulation (2-3 s) periods. We first evaluate the brain state from a regional perspective based on the properties of the frequency domain. We subtract the corresponding mean value from each excitatory time series and apply Welch’s method with a window length of 0.5 s with 50% overlap to estimate the power spectrum density of each area. The peak frequency is given by

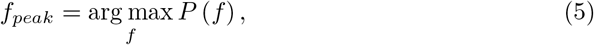

Thus, the frequency at which the regional power reaches its maximum value is the measure of interest. In this work, due to the general low-peak frequency of brain areas before the perturbation, the peak frequency of 2-3 s captures the impact of local stimulation.

We also consider the brain state from a network viewpoint based on functional connectivity, which is derived by calculating the maximum normalized cross-correlation [100, 101] between time series with a time window of 1 s and a maximum lag of 250 ms. The stimulation effects are quantified as the difference in the dynamic behaviors in the before-(1-2 s) and during-stimulation (2-3 s) periods, namely, the functional and structural effects [37]. The functional effect (*fe*), which reflects the influence of local brain regions on the interregional coupling configuration, is calculated as

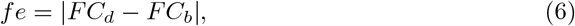

where *FC*_*d*_ and *FC*_*b*_ are the functional connectivity matrices during and before stimulation, respectively, and || represents the average of the absolute values of elements in the upper triangle of the matrix. The structural effect (*se*), which reflects how local changes in regional activity affect the structural constraints on brain dynamics, is given by

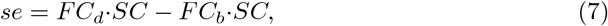

where *SC* is the structural connectivity matrix and · indicates calculating the Pearson correlation coefficient between two matrices.

In addition to these basic metrics, various integrated measures were used in this study to characterize the performance of the brain. These measures are listed in Table 2 for ease of review.

**Table 2.**
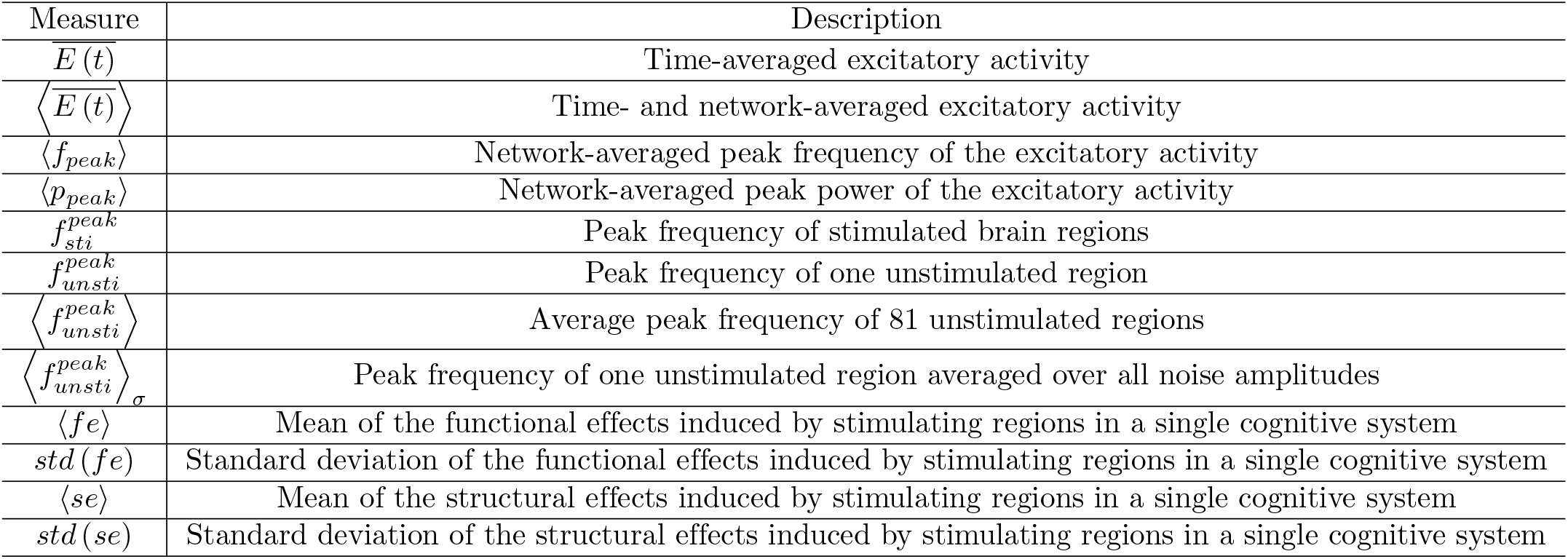
Summary of integrated measures.

## Supporting information

**S1 Fig. Distribution of the peak frequency of the excitatory activity in all brain regions under different noise amplitudes given that** *c* = 0.1. (A) Noise amplitude = 10^*−*7^. (B) Noise amplitude = 10^*−*5^. (C) Noise amplitude = 10^*−*3^. (D) Noise amplitude = 10^*−*2^. Panels (A-C) show similar behaviors, remarkably different from that of panel (D). These results are comparable to Fig. 2C.

**S2 Fig. Distribution of the peak power of the excitatory activity in all brain regions under different noise amplitudes given that** *c* = 0.1. (A) Noise amplitude = 10^*−*7^. (B) Noise amplitude = 10^*−*5^. (C) Noise amplitude = 10^*−*3^. (D) Noise amplitude = 10^*−*2^. Noise amplitude increases the peak power in all brain regions, similar to results in Fig. 2D.

**S3 Fig. Robustness of the effect of noise amplitude (x-axis) on the peak frequency of unstimulated brain regions** 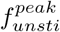 **(y-axis) at various stimulation sites**. (A) L-Pars Orbitalis (small degree). (B) R-Superior Frontal (moderate degree). (C) L-Caudate (large degree).

**S4 Fig. Robustness of the impact of local perturbations on the time series and power spectra of unstimulated brain regions**. The upper panels indicate the time series of two unstimulated brain regions at different noise amplitudes when stimulating the R-Lateral Orbitofrontal region. The lower panels indicate the power spectra before (blue) and after (orange) stimulation in the corresponding condition. (A) L-Caudate, noise amplitude = 10^*−*7^. (B) L-Caudate, noise amplitude = 10^*−*5^. (C) L-Caudate, noise amplitude = 10^*−*3^. (D) L-Pars Orbitalis, noise amplitude = 10^*−*7^. (E) L-Pars Orbitalis, noise amplitude = 10^*−*5^. (F) L-Pars Orbitalis, noise amplitude = 10^*−*3^.

**S5 Fig. Oscillations before stimulation have little impact on the similarity between the peak frequency averaged across various noise amplitudes and the structural connectivity**. (A) The peak frequency 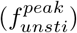 of unstimulated brain regions (y-axis) under different stimulated brain regions (x-axis) at a large noise amplitude (10^*−*2^, corresponding to oscillations before stimulation). The diagonal elements are set to 0. (B) The peak frequency 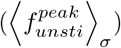of unstimulated brain regions (y-axis) under different stimulation sites (x-axis) averaged across various noise amplitudes that do not induce oscillations before the perturbation. (C) The positive Spearman correlation (*r* = 0.95, *p <* 0.01) between the matrix in (B) and the structural network is similar to that in Fig. 4(e).

**S6 Fig. Examples of functional connectivity changes caused by stimulating different brain regions in oscillatory states before perturbation under a large noise amplitude (**10^*−*2^**), showing similar results to Fig. 5**. (A) R-Lateral Orbitofrontal. (B) R-Hippocampus. (C) L-Accumbens.

**S7 Fig. Snapshots of functional effects under other noise amplitudes for one realization**. (A) Noise amplitude = 10^*−*8^, Pearson’s *r* = 0.67, *p <* 0.01. (B) Noise amplitude = 10^*−*6^, Pearson’s *r* = 0.79, *p <* 0.01. (C) Noise amplitude = 10^*−*4^, Pearson’s *r* = 0.94, *p <* 0.01. (D) Noise amplitude = 10^*−*2^, Pearson’s *r* = 0.16, *p* = 0.1447. The gray lines represent the linear fits of data points estimated by ordinary least squares. These results are in line with the trend shown in Fig. 6C.

**S8 Fig. Snapshots of structural effects under other noise amplitudes for one realization**. (A) Noise amplitude = 10^*−*8^, Pearson’s *r* = −0.56, *p <* 0.01. (B) Noise amplitude = 10^*−*6^, Pearson’s *r* = −0.39, *p <* 0.01. (C) Noise amplitude = 10^*−*4^, Pearson’s *r* = 0.08, *p <* 0.4991. (D) Noise amplitude = 10^*−*2^, Pearson’s *r* = 0.32, *p <* 0.01. The gray lines represent the linear fits of data points estimated by ordinary least squares. These results are comparable to the trend shown in Fig. 7C.

**S9 Fig. Locations of cognitive systems in terms of functional and structural effects under other noise amplitudes**. (A) Noise amplitude = 10^*−*8^. (B) Noise amplitude = 10^*−*7^. (C) Noise amplitude = 10^*−*5^. (D) Noise amplitude = 10^*−*4^. (E) Noise amplitude = 10^*−*2^. Note that stimulated brain regions are grouped into 4 cognitive systems with different colors. The colored areas represent the convex hulls of data points in the systems. The points reflect the measures averaged over 30 realizations.

## Notes

### Competing Interest Statement

The authors have declared no competing interest.

